## Supporting for "Wisent genome assembly uncovers extended runs of homozygosity and a large deletion that inactivates the thyroid hormone responsive gene": Supplementary Figures.docx

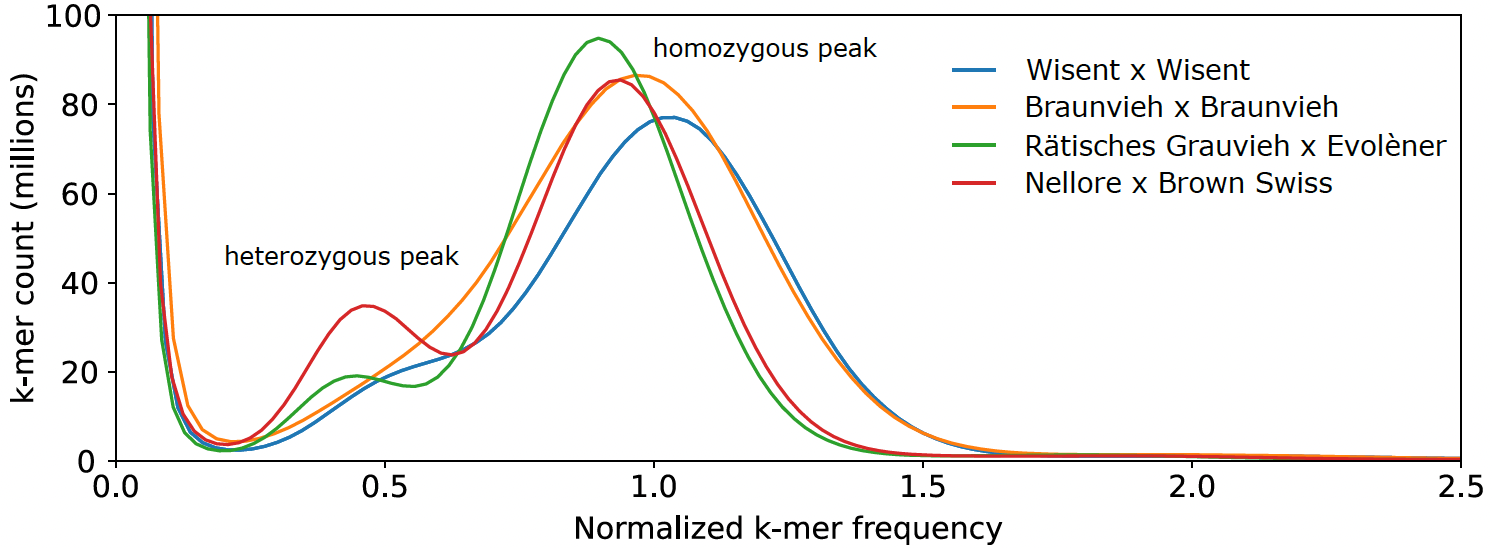

**Supplementary Figure 1: Distinct k-mer distribution for four samples.** Although the wisent sample has lengthy runs of homozygosity, its heterozygous peak is slightly more defined than in the Braunvieh x Braunvieh sample.

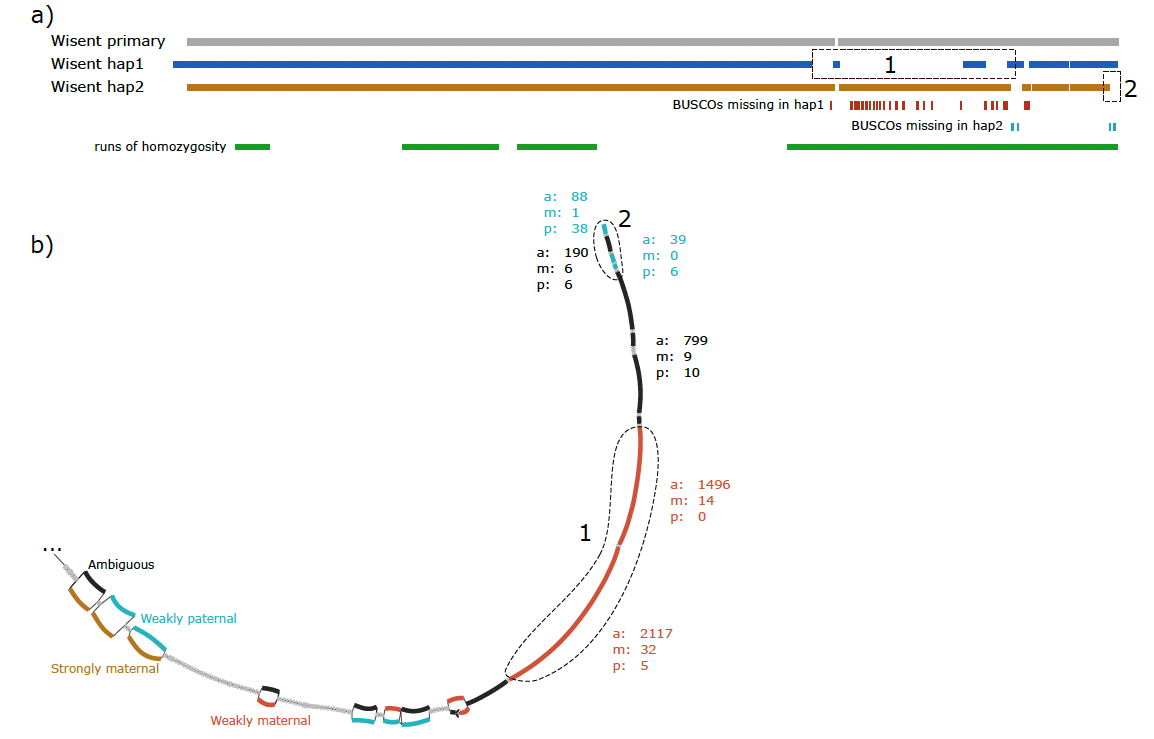

**Supplementary Figure 2: Missing BUSCO genes are located in regions entirely missing from one haplotype.** (a) Pangenome synteny of the primary and haplotype-resolved wisent assemblies on chromosome 17. Missing BUSCOs are shown below, corresponding to sequence missing from one of the two haplotype-resolved assemblies. (1) is an approximately 15 Mb of sequence containing 24 BUSCOs missing from haplotype 1 but present in haplotype 2 and the primary assembly while (2) is approximately 1 Mb and contains 2 BUSCOs missing from haplotype 2. Runs of homozygosity larger than 2 Mb and fewer than 2 heterozygous variants per 10 Kb are shown. (b) An annotated region of the hifiasm primary unitig graph on chromosome 17 from 40 Mb to the end. Large unitigs can be assigned as Ambiguous (black) if there is no imbalance in parental haplotype tagging. Unitigs could also have primarily maternal tagged reads (brown), primarily ambiguous but biased towards maternal (orange), or primarily ambiguous but biased towards paternal (blue). This region did not contain any strongly paternal tagged unitigs. The major haplotype-resolved errors from regions (1) and (2) were weakly maternal or paternal respectively, located within long runs of homozygosity. The sequences appear to be homozygous but were incorrectly assigned to only one haplotype during haplotype-phasing of this graph.

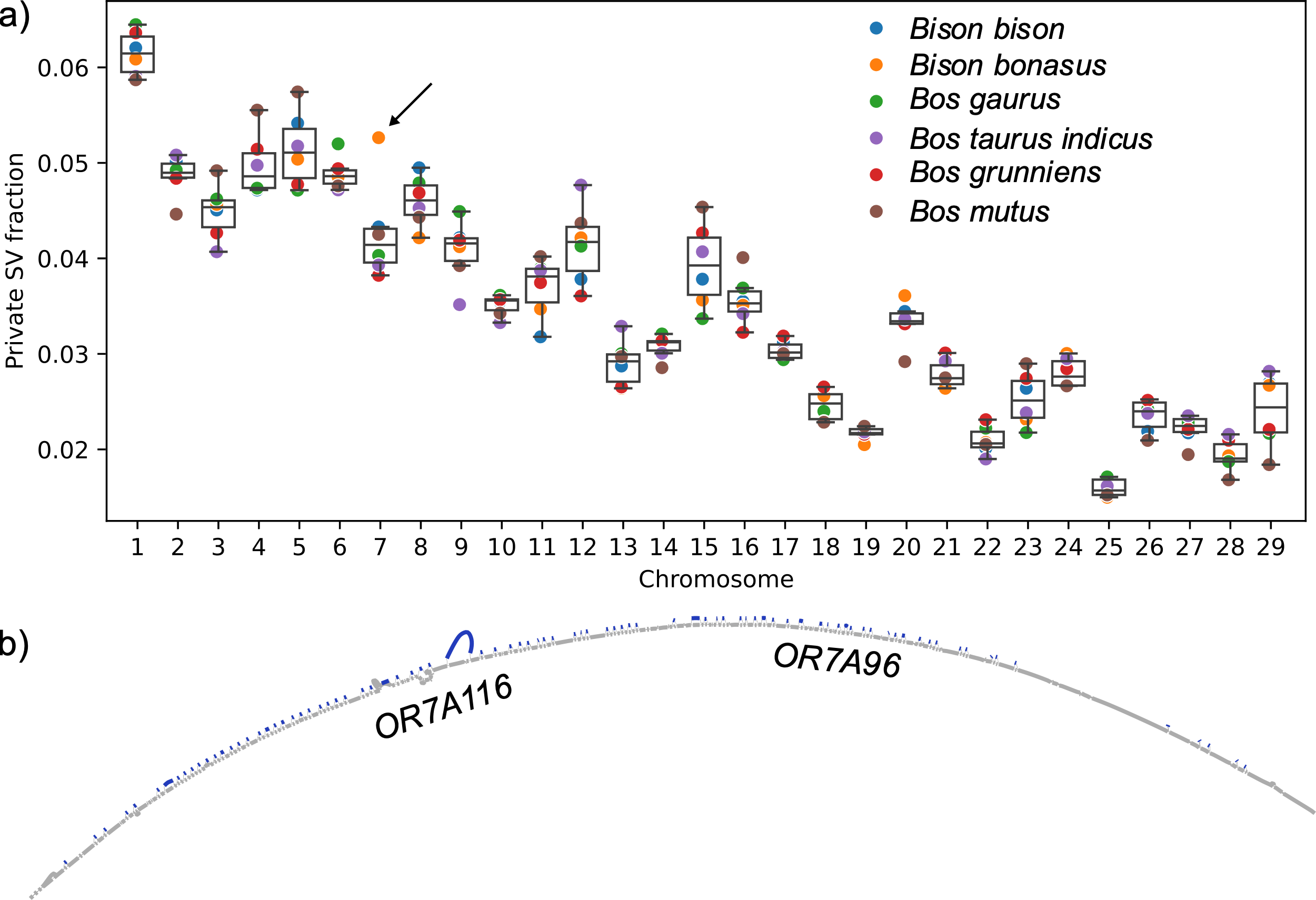

**Supplementary Figure 3: Excess of private structural variants to wisent on chromosome 7.** (a) Chromosome 7 was a substantial outlier when considering the number of SVs private to wisent compared to the total number of SVs per chromosome and was not a pattern observed in any other sample. (b) Most private SVs to wisent were clustered between 10 and 10.6 Mb on chromosome 7, a region containing 12 annotated protein-coding genes. Two of these genes are explicitly identified as part of the olfactory receptor family 7 subfamily A (OR7A), while many of the remaining genes have orthologous relationships to the OR7A subfamily.

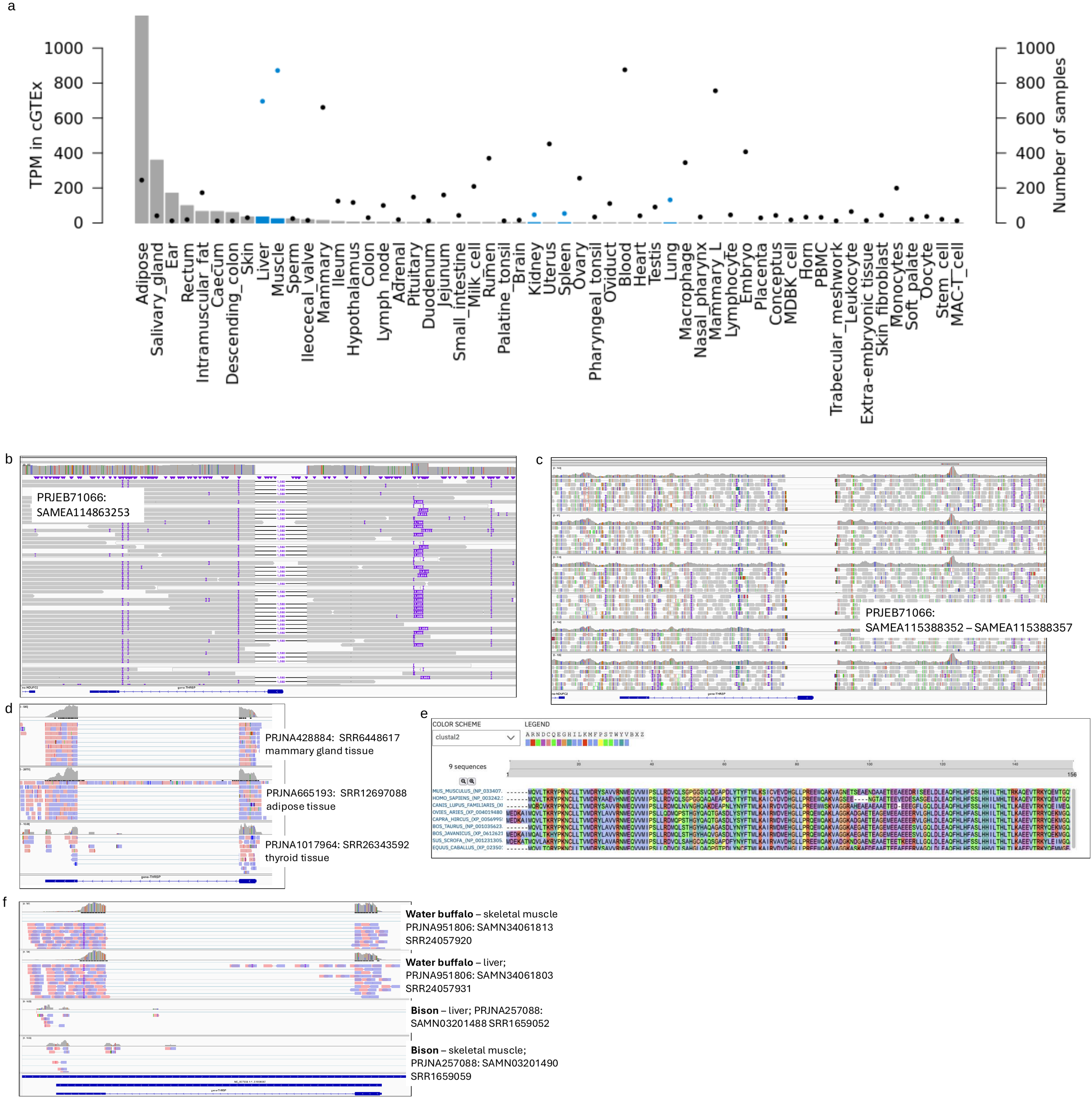

**Supplementary Figure 4: Validation of a large deletion uncovered by the pangenome analysis.** (a) Expression of *THRSP* in 55 tissues from the cattle GTEx dataset. The bars represent TPM values, and the black dots represent the number of samples per tissue. The y axis is truncated at 1000. Blue colour represents tissues for which transcriptome data are also available for the 3 years old bison cow. (b) Alignment of the wisent F1 HiFi reads against ARS-UCD1.2 confirm deletion of 1,580 bp sequence encompassing the first coding exon of THRSP encoding thyroid hormone responsive protein. (c) Alignment of six short-read sequenced wisent genomes support the deletion. (d) Alignment of RNA sequencing data from three cattle tissues (mammary gland, adipose, thyroid) confirm expression of the first THRSP exon. (e) Multi-species alignment of the THRSP protein sequence. (f) Alignment of RNA sequencing data from skeletal muscle and liver tissue from water buffalo and bison. Both exons are expressed in water buffalo while neither of the exons is expressed in bison.
